# Uncertainty during visual search: Insights from a computational model and behavioral experiment in natural stimuli

**DOI:** 10.1101/2025.01.14.632909

**Authors:** Gaston Bujia, Gonzalo Ruarte, Melanie Sclar, Guillermo Solovey, Juan E. Kamienkowski

## Abstract

Visual search, driven by bottom-up and top-down processes, offers a unique framework for investigating decision-making. This study examines individuals’ awareness of their own visual search by combining computational modeling with behavioral experiments. Fifty-seven participants performed a classical visual search task in which the goal was to find an object in a natural scene. Crucially, in some trials, the search was interrupted by clearing the screen before the gaze reached the target object. Participants had to report their best guess of the target’s location and the uncertainty on their response. We show that a modified version of the Entropy-Limit Minimization (ELM) model captures scanpaths and perceived target locations, while also revealing that uncertainty is influenced by scanpath length, the distance between the perceived and true target location, and the entropy of decision maps. These findings highlight the model’s capacity to reflect cognitive processes underlying response selection and uncertainty judgment.

## 1 Introduction

Visual search is a sequential decision-making task that requires integration of bottom-up and top-down information involving multiple cognitive processes [Bujia et al., 2022, Eckstein, 2011, Treisman and Gelade, 1980]. Bottom-up mechanisms are stimulus-driven, relying on the inherent salience of visual features to guide attention toward certain regions in the visual field. These features include brightness, color, or movement, which naturally attract attention. Conversely, top-down mechanisms are goal-driven, operating under voluntary control to align attention with specific objectives or tasks. Attention in visual search is driven by both voluntary, goal-driven mechanisms and involuntary, stimulus-driven responses to perceptual properties [Eimer, 2014]. As visual search is far from random, progress is made as participants explore the scene. This raises critical questions: Are individuals aware that they are close to finding the target object? Does individuals’ uncertainty about the target location reflect the objective information acquired during search? In this work, we use a naturalistic visual search task coupled with computational models to address these questions.

Computational models have proven to be a powerful tool for investigating complex human behavior and its interplay with cognition processes, ranging from modeling learning and inference [Tenenbaum et al., 2006, Friston, 2005], to exploring the nature of visual decision processes [Sigman and Dehaene, 2005, Wolfe, 2021] or the construction of confidence and uncertainty [Zylberberg et al., 2012, Pouget et al., 2016]. In visual search, a wide variety of theoretical frameworks have been proposed for modeling scanpath behavior in different scenarios and for diverse purposes [Sclar et al., 2020, Bujia et al., 2022, Najemnik and Geisler, 2005, Rashidi et al., 2023, Zelinsky, 2008, Zhang et al., 2018]. Najemnik and Geisler [2005] proposed a pure Bayesian approach for modeling the fixation selection during visual search, meanwhile Zhang et al. [2018] implemented a strategy solely based on pre-trained deep neural networks. Other works proposed a combination of both approaches, showing models that can accurately predict human scanpaths during visual search tasks in different natural scenarios [Travi et al., 2022]. This blend of computational approaches enriches our understanding of visual selection mechanisms.

However, with a few exceptions, the question of how subjective uncertainty about a target location evolves in time during a visual search task remains relatively unexplored. Specifically, understanding the extent to which individuals are conscious of their computational processes during the task and whether models that capture eye movements can bridge this gap between them and the intricate cognitive process that lies under them. Self-reports have been used to probe subjective experiences in visual search; Kotowicz and collaborators [Kotowicz et al., 2010] employed a visual search paradigm with random early termination and found that confidence, but not necessarily accuracy in task performance, increases with the time spent within a fixation. More recently, Adams and Gaspelin [2021] conducted a singleton search paradigm [Theeuwes, 1992] and demonstrated that participants exhibited a clear internal level of introspective awareness of attentional capture. Specifically, participants were able to monitor attentional shifts even in the presence of distractors. Expanding on these findings, Mazor and Fleming [2022], tested several hypotheses about efficient search termination in a feature search task, collecting self-reports on task-related knowledge. They showed that while some knowledge of search difficulty was often accessible for reporting, this was not a necessary condition for efficient search termination. However, all these previous works were conducted in controlled, artificial paradigms.

To delve deeper into the complexities of behavior and the need for strong analytical tools, researchers have turned to advanced computational methods such as entropy measures and persistent homology diagrams. Persistent homology, rooted in topological data analysis, is adept at identifying geometrical features across different scales of data, uncovering complex patterns that traditional analyses might overlook [Chazal and Michel, 2021, Chintakunta et al., 2015, Merelli et al., 2015, Turkes et al., 2022]. It has been shown to be effective in quantifying information and finding structures in different problems as feature extractors or descriptors. For instance, Garner et al. use persistence diagrams to describe the underlying structure of the spiking activity of grid cells in the medial entorhinal cortex [Gardner et al., 2022], and Guo et al. [2022] developed a multi-scale brain network modeling method using persistence features for electroencephalography to explore brain functional networks in schizophrenia patients. The combination of these measures with Entropy provides a strong framework to assess the information entangled in structured data from complex decision-making tasks like visual search [Ribeiro et al., 2012].

In this work, we seek to answer: (1) To what extent do participants’ uncertainty reports align with their search performance and response behavior? and, (2) Can computational models predict and explain the subjective uncertainty experienced by participants during naturalistic visual search? To address these questions, we conduct and extend an overt visual search experiment in natural scenes. Particularly, we evaluate the awareness of the human decision-making process by asking participants to provide both the perceived target location (clicking on where they believed the target was) and an uncertainty response (drawing a circle around the target). We analyze the experimental results and compare them with an adaptation of the Entropy-Limit Minimization model [Najemnik and Geisler, 2009], the neural network ELM (nnELM. We then compare human and model predictions of the target location, and whether the model’s internal states can explain individuals’ uncertainty about the target location.

## 2 Methods

### 2.1 Participants, paradigm, and data acquisition

In the present work we use the same data collected in Sclar et al. [2020] (see also Travi et al. [2022]) with the difference that in these previous publications, we focused on the modeling of the eye movements, and now we extend the model to the responses after the search is finished. Eye movements were recorded with an Eye Link 1000 (SR Research, Ontario, Canada) monocular at 1000 Hz. Saccades and fixations were parsed online with the native algorithm. The experiment was programmed using PsychToolbox and Eye Link libraries in MATLAB [Brainard, 1997] in a 19-inch Samsung SyncMaster 997 MB monitor (refresh rate = 60 Hz, resolution = 960 *×* 1280). All the experiments described here were reviewed and approved by the ethics committee: “Comite de Ética del Centro de Educación Médica e Investigaciones Clínicas ‘Norberto Quirno’ (CEMIC)” (qualified by the Department of Health and Human Services (HHS, USA): IRb00001745—IORG 0001315; Protocol number 435).

Fifty-seven subjects (34 male, 23 female; age 25.1 *±* 5.9 years old) had to search for an object in a crowded grayscale indoor scene. Each trial started when the target was presented in the center of the screen. After 3 seconds, the target was replaced by an initial fixation dot at a pseudo-random position at least 300 pixels away from the actual target position in the image. This was done to avoid starting the search too close to the target (Fig. 1A). The initial position was the same for a given image and all participants. The search image appeared after the participant fixated the initial dot. The image was presented at a 768 *×* 1024 resolution (subtending 28.8 degrees of visual angle). The search period finished when the participant fixated on the target or after *N* saccades, with an extra 200ms to allow observers to process information of this last fixation [Kotowicz et al., 2010]. The maximum number of saccades allowed (*N*) were 2 (13.4% of the trials), 4 (14.9%), 8 (29.9%) or 12 (41.8%). These values were randomized for each participant, independent of the image. Each trial was considered successful (the target was effectively found) if any fixations landed on the target’s box subtending approximately 1.25° degrees of visual angle or at much 72 pixels of distance from the center of the box. All participants completed a total of 134 trials with different targets and backgrounds in three blocks.

**Figure 1:**
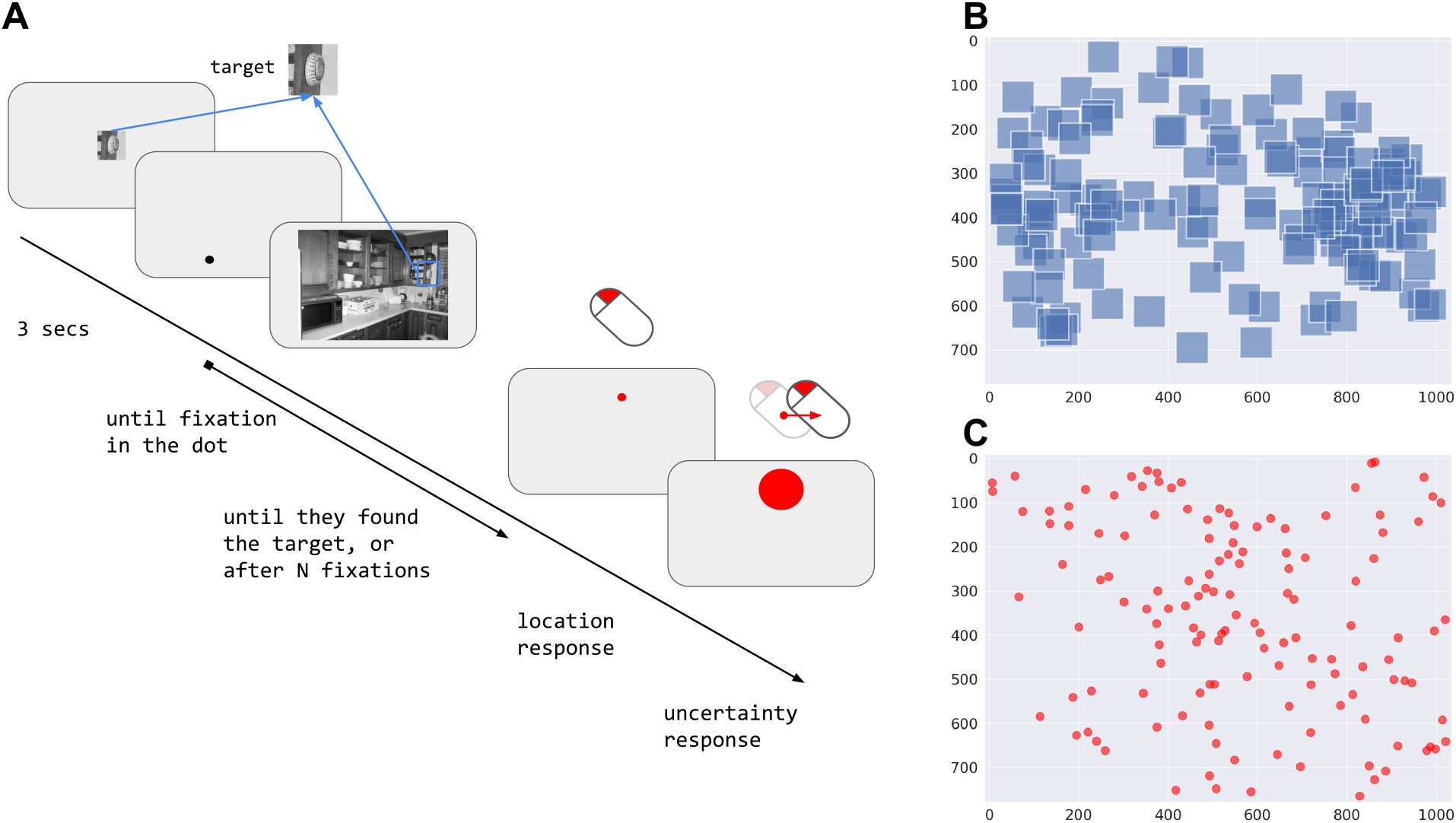
Experimental design. **A**. Timeline. The target was displayed at the center of the screen on a gray background at twice its original size for 3 seconds. After that, a dot was presented until fixated, starting the trial. The trial finished when the target was found or after the participant performed N fixations. After the trial ended, the participant reported the target location by selecting a point with the left mouse button and then reported their uncertainty by determining the radius length using the mouse buttons. **B**. Spatial distribution of targets (blue squares), and **C**. Spatial distribution of first fixations (red dots).

After the search finished, a screen with only the frame of the image and a mouse pointer (a small black dot) was displayed on the screen. The participant was asked to indicate where he/she thinks the target is located. They had a black circle of variable center and radius and were asked to choose the center and radius such that the circle covered the position where he/she thought the target was located. First, the participants selected a position with the mouse, the center of the circle (location response or response), and second, they indicated the uncertainty of their decision by increasing or decreasing the radius of the circle with the mouse buttons (uncertainty response or response size). The minimum size of the radius was set at 3 pixels. Each response was considered successful if it landed at a distance less than or equal to 72 pixels from the center of the box.

### 2.2 Neural Network Entropy-Limit Minimization Model

In this work, we chose to use an adaptation of the visual search model proposed by Najemnik and Geisler: the Entropy Limit Minimization model or ELM [Najemnik and Geisler, 2009, Zhou and Yu, 2021, Rashidi et al., 2023]. The original model showed near-optimal performance in visual search on artificial stimuli and produced human-like eye movement statistics. The objective of the ELM is to minimize the uncertainty of the posterior probability of the target location by maximizing the information gain on the next saccade. To do so, it takes into account the visibility map and the current fixation location, then it considers all possible target locations on a grid (we used a cell grid size of 32 pixels spanning a 24 *×* 32 grid) and chooses the location that minimizes the expected Shannon entropy of the posterior probability distribution. To work with natural stimuli, we first adapted the ELM to use a saliency map of the search image computed with DeepGaze II [Kummerer et al., 2017] (Fig. S1) as the prior probability of the target location. Then, ELM estimates the posterior probability of finding the target on the i-th location at the (T+1)-th fixation *p*_*i*_(*T* + 1) (eq. 1) as the combination of the prior probability in the previous location and the squared visibility or detectability (*d*) weighted by the template response (*W*).

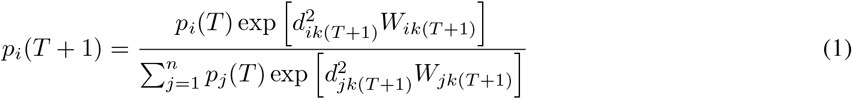

This can be rewritten more generally as:

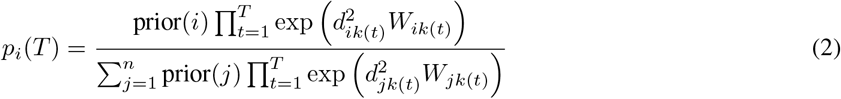

The visibility map *d*_*ik*(*T*)_ represents how visible the target is on the location *i* when the observer is located at *k*(*T*) (it is quantified as a fixed 2D Gaussian centered on the current fixation). Meanwhile, *W*_*ik*(*T*)_ represents the template response that measures the information collected from the location *i* given the observer location *k*(*T*). This template response was originally a random sample drawn from a Gaussian distribution *W*_*ik*(*T* +1)_ ∼*N*(*µ*_*ik*(*T*)_, *σ*_*ik*(*T*)_) that depended only on the presence or absence of the target and the inverse of the visibility map (eq. 3). Here, we incorporate the notion of similarity between the target and each image location (i.e. similarity map, see below)

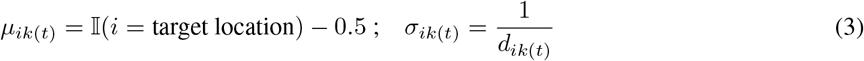

To calculate the next fixation, the ELM estimates the expected reduction of entropy as the current posterior probability distribution across the possible target locations weighted by the visibility (eq. 4):

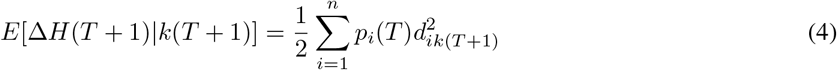

Given a fixation order *T*, eq. 4 inherently defines a mapping between all possible target locations on the scene and the expected information gain for each of them. This map is used later for calculating the human scanpath metrics (see Metrics section) and we call it decision map. Finally, the next fixation *k* is selected as the one that maximizes the information gain obtained from selecting that location *k*_*opt*_(*T* + 1) (eq. 5):

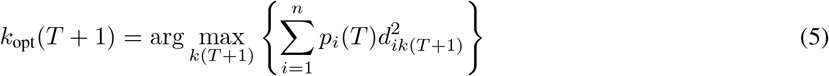

The original ELM approach was entirely proposed, and tested in artificial stimuli where subjects had to search for a Gabor patch embedded in 1/f noise. Further adaptations were proposed for modeling other characteristics of the scanpaths like the fixation duration [Zhou and Yu, 2021] or incorporating color [Rashidi et al., 2023]. In the case of natural stimuli, we followed the original adaptation proposal made in Bujia et al. [2022], Sclar et al. [2020], Travi et al. [2022], the idea is to introduce a notion of similarity map *ϕ* into the template response via modifications into the parameters of the Gaussian distribution. Previously, it was performed with a mean value that combined the visibility and the similarity calculated with cross-correlation or structural similarity between the target template and the i-th location of the image (*ϕ*_*i*_) [Sclar et al., 2020, Bujia et al., 2022] and, later on, with the attention map from a deep neural network [Zhang et al., 2018]. We will use the output of the IVSN network [Zhang et al., 2018] at location *i* (eq. 7) in combination with the proposal for the template response (eq. 6):

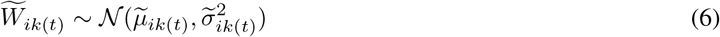

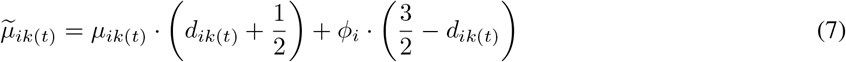

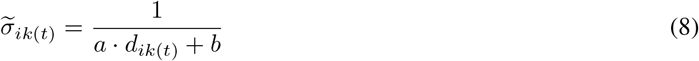

Where, *a* and *b* in eq. 8 represent terms for avoiding dividing by zero. By performing this substitution, the resulting model can capture object invariance while maintaining good performance on template matching.

### 2.3 Scanpath Metrics

To compare stepwise the decision-making processes of the model with those of humans, we utilized the Human Scanpath Prediction (HSP) metrics proposed in the ViSiONS benchmark [Travi et al., 2022]. This metric allowed us to compare the scanpath at a fixation level, for a given participant’s scanpath, we calculated how well each model predicts the next human fixation based on the scanpath history. The fundamental concept of this method is to compel the model to follow the participant’s scanpath, disregarding its own predictions. Generally, this metric evaluates for each fixation a model’s conditional priority map (a priority map informed by the model’s internal state, the posteriors, and the previous subject’s fixations) and compares it against the participant’s fixation [Kümmerer and Bethge, 2021]. Using the conditional priority map as a classifier as the ground truth allows for the computation of well-established metrics such as AUC or the Normalized Scanpath Saliency (NSS) [Bylinskii et al., 2019, Peters et al., 2005], and Information Gain or Log Likelihood [Kümmerer et al., 2015] relative to the center bias and uniform baselines. Higher values are better for all these metrics. For the ELM specifically, we chose only two to analyze deeper, the AUC and IG, components (Fig. 4), and we used the decision map defined by equation 4 as the conditional priority map. We also included in the benchmark’s public repository the comparison of the results of the nnELM model on the estimation of the eye movements in the present dataset with previous models (Github’s repository). For more details see Travi et al. [2022].

### 2.4 Entropy

We employed multiple entropy measures to comprehensively assess the information content of the decision maps of ELM. We proposed to use the classical Shannon Entropy but also the Fuzzy 2D Entropy (**Fuzz2D**) [Hilal and Humeau-Heurtier, 2019], Distributional 2D Entropy (**Dist2D**) [Azami et al., 2017], Sampling 2D Entropy (**Samp2D**) [Silva et al., 2016] and Permutation 2D (**Perm2D**) [Flood and Grimm, 2021, Ribeiro et al., 2012]. Except for the Shannon Entropy, the rest of the proposals here considered are two-dimensional extensions of the entropy. These were majorly designed to capture the complexity of geometrical patterns in images. Dist2D, Samp2D, and Fuzz2D were proposed to separate textures on images for medical uses [Hilal and Humeau-Heurtier, 2019], while Perm2D was developed to study time variations on images. This approach of considering other entropies allows us to test several possible ways decision maps may encode relevant information to the task. We used the implementation of the Entropy Hub package in Python [Flood and Grimm, 2021].

### 2.5 Persistence Homology Metrics

We incorporated features from topological data analysis and persistence homology to further quantify the spatial information encoded on decision maps by the ELM. We computed the 0-dimensional homologic persistence diagram or barcode corresponding to the upper-level set filtration [Bauer, 2021, Chazal and Michel, 2021, Ghrist, 2008, Tralie et al., 2018]. For each decision map, we calculated the persistence barcode and extracted two different features: the number of bars (denoted as **H0**), which serves as a proxy of the connected components of maps, and the persistence entropy [Myers et al., 2019], i.e. the entropy of the lengths of the bars in the barcode (denoted as **PE**). Also, we considered other two topological features: we filtered the barcode by erasing the bars that were shorter than the mean bar length in the barcodes of the rest of the decision maps (i.e. all maps related to any other fixation that was not the last fixation or the response). We computed the number of connected components (**H0f**) and the persistence entropy (**PEf**) for those barcodes. For more details, see supplementary material. We calculated these features using the Python implementation Ripser [Tralie et al., 2018].

### 2.6 Statistics

To evaluate the relationship between response and overall behavior, we grouped the trials into four categories based on whether the target was found during the search and whether it was selected with the participant’s subjective response (see Fig. 1A). Thus, each trial was classified according to two criteria: (1) if subjects saw the target during the search, it was marked as **target found online** (**TFO**) and as **∼TFO** otherwise; and (2) if subjects identified the target with their location response, it was marked as **target found response** (**TFR**), and as **∼TFR** otherwise (see Table 1; Fig. 3C). A response was considered as TFR based only on the location response whether it was at a distance of up to 72 pixels from the center of the target bounding box. Trials classified as TFO & ∼TFR were discarded due to their low prevalence (less than 6% of trials) and because they were considered errors in the responses of the participants.

**Table 1:**
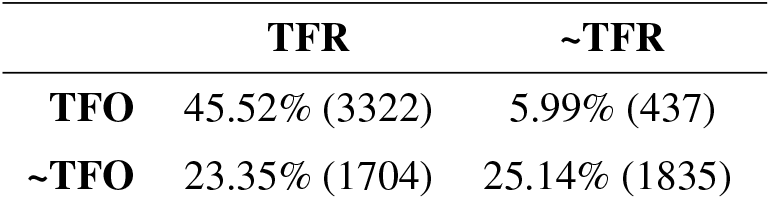
Classification of trials according to the search and response results. TFO: target found online (during the search) and TFR: target found response (with the manual response after the search).

We used linear mixed-effects (LMMs) models to quantify relationships between the different variables considered using a random effect for each subject and each image. For the LMMs, regression coefficients (*β*s), t-values (*t* = *β/SE*), and 95% confidence intervals were reported, and SE can be inputted from those intervals. There is no definitive definition of “degrees of freedom” for LMMs, so precise p-values cannot be calculated [Arnqvist, 2020]. We considered an effect significant if |*t*| = |*β/SE*| *>* 1.96. Transformations performed to the considered variables are also reported on each corresponding table. Model fitting and the significance of fixed effects were assessed using the lme4 package and the confint function. We also reported the conditional-*R*^2^ which represents the variance explained by the model, including both fixed and random effects. The conditional-*R*^2^ and its components varied across the different models, as the relative contributions of fixed and random effects to the total variance depend on the model structure [Nakagawa et al., 2017]. It was estimated for each model using the Multi-Model Inference package in R [Bartoń, 2024], Tables S1, S2, S3 and S4).

## 3 Results

### 3.1 Behavioral results

We explored the overall performance of the participants’ responses by looking first at the location responses, i.e. where exactly they expect the target to be (Fig. 2); and second, at the uncertainty responses, i.e. the confidence of the responses measured by the estimated area in which they expect the target could be (Fig. 3). For the objective responses, we measured two distances: the Euclidean distance between the center of the target and the response and, the distance between the response and the last fixation of the scanpath. We grouped trials according to two conditions: the saccadic threshold or maximum allowed fixations (2, 4, 8, or 12) and whether the target was found or not. As expected, participants who did not find the target, responded on average further than the ones who did find it for all thresholds (Fig. 2A, C, Table S1, *t*(*MaxFix*) = −11.42). Moreover, as more saccades were allowed, the distance from the response to the target was reduced on average in not-found trails (Fig. 2A, C). On the contrary, when subjects found the target, they were consistent in their response independently of the number of saccades allowed (Fig. 2A, C, Table S1, *t*(*TF*) = −21.36; *t*(*MaxFix* : *TF*) = 4.76). When comparing the distribution of distances between the response and last fixation we also observed that the target-found trials showed a smaller distance between the response and the last fixation than not-found (Table S1, *t*(*MaxFix*) = −9.00; *t*(*TF*) = −40.25; *t*(*MaxFix* : *TF*) = 5.97; Fig. 2B, C). Nevertheless, the distance seems to be less affected by the number of saccades allowed in any of the target-found conditions (Fig. 2B, C).

**Figure 2:**
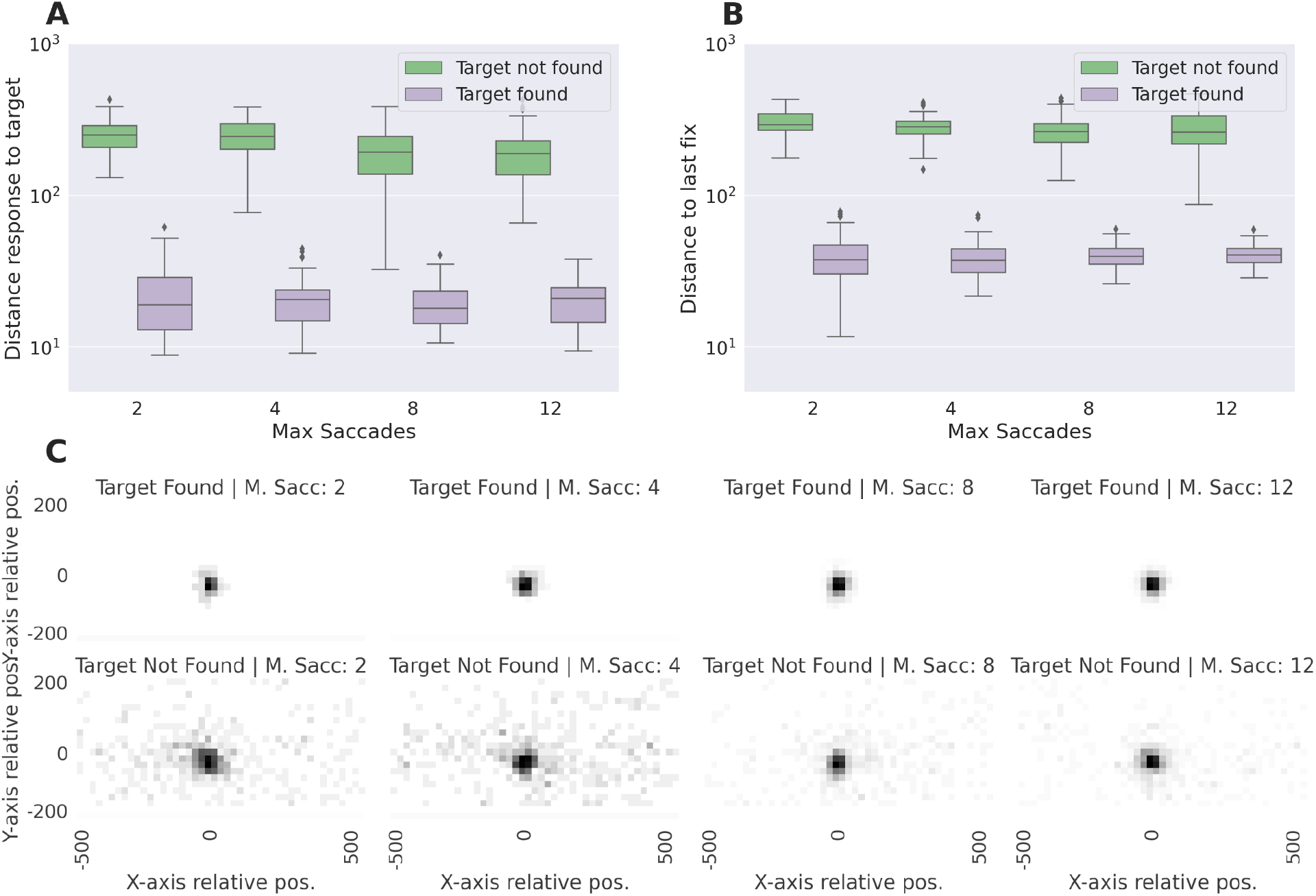
Objective response. **A**. Euclidean distance from the response to the target, **B**. Euclidean distance from the response to the last fixation, and, C. spatial distribution of responses relative to the target at the center, all as a function of the found condition of the trial (found or not found) and the maximum number of saccades allowed. (Each point is subject, error bars correspond to the standard deviation). In the rightmost of panel **C**., the color bar represents the density mapping for all the heatmaps in that row.

**Figure 3:**
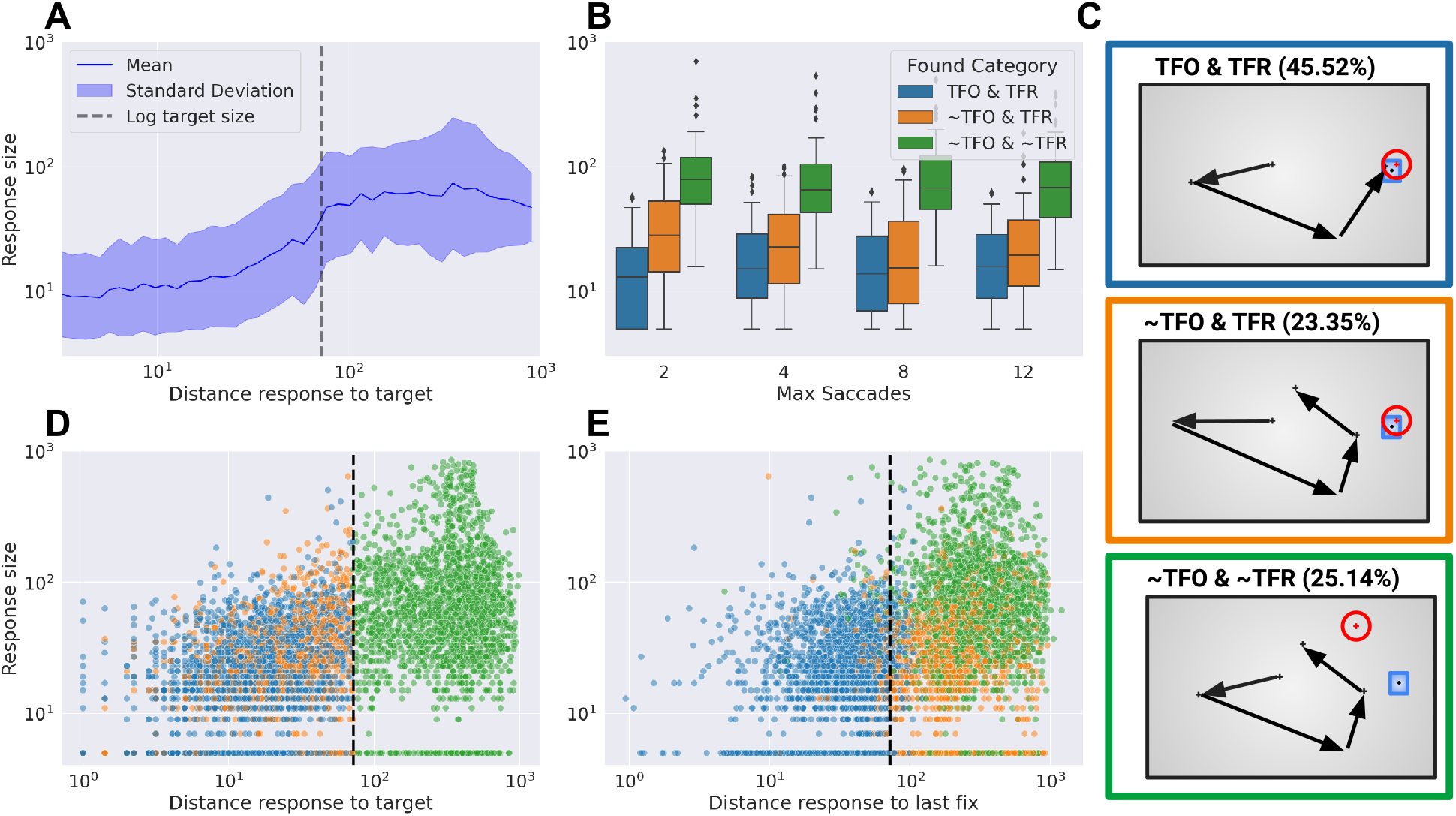
Uncertainty response. **A**. Logarithm of the uncertainty (or response size) as a function of the distance of the response to the target for all responses considered. **B**. Uncertainty, measured as the size of the response, as a function of the maximum of saccades allowed and grouped by the trial category. **C**. Scheme of the trial categories. **D**. Distribution of distance from response to target and the response size for each trial category and distribution of distance from last subject fixation to the response and the response size (**E**.). The dashed line represents the threshold for considering a response as target found.

When focusing on the uncertainty report, results showed that the response size stabilized (plateaued) as the error, measured by the distance from the response to the target, increased beyond a certain threshold close to the target size (Fig. 3A). Our study also revealed a significant decrease in uncertainty when the target was found (Fig. 3B; Table S1, *t*(*TF*) = −40.25). This was more evident when disaggregating the implicit behavior of the not found condition into the explicit behavior for the response found condition (∼TFO & TFR/∼TFO & TFR vs. ∼TFO & ∼TFR/∼TFO & ∼TFR, orange vs green respectively in Fig. 3B-D). Those effects were positive (increased uncertainty), both statistically significant (Fig. 3B-D; Table S1, *t*(∼*TFO&TFR*) = 7.28, *t*(∼*TFO*&∼*TFR*) = 23.90), for the first case was weaker than the second case, indicating a less pronounced uncertainty when participants explicitly identified the target after an initially implicit failure, compared to when the target remained undetected. Additionally, the results showed that the number of allowed fixations (MaxFix) played a significant role in reducing uncertainty when not accounting for the response condition (Fig. 3B,C; Table S1, *t*(*MaxFix*) = −8.53). Nonetheless, when incorporating the response condition, that main effect lost its significance (Table S1, *t*(*MaxFix*) = 0.17) but the interactions with the TFR found condition resulted significant (Table S1, *t*(*MaxFix* : *TF* − *NF*) = −3.91, *t*(*MaxFix* : ∼*TFO*&∼*TFR*) = − 1.76). These findings, depicted in Figure 3, highlight that increased uncertainty in the absence of target identification was more influenced by the actual outcome/performance rather than the quantity of information gathered during the search process.

To further explore the relationship between the uncertainty and both distances (response-target, response-last fixation), we evaluated the Spearman correlation between both variables and the response size (Fig. 3D). Interestingly, uncertainty was positively correlated with the distance of the response to the location of the target and with the last fixation (target response distance *r* = 0.56, last fixation response distance: *r* = 0.45; both p-value < 10e-3, N = 6883) in general. Moreover, both behaviors were shown to have a stronger significant relation with uncertainty in cases where the participants could not find the target only but effectively responded correctly “∼TFO & TFR” (target response distance *r* = 0.31; last fixation response distance: *r* = 0.22; both p-value < 10e-3, N = 1507), but in cases where the subjects answered wrongly the distance to the target lost its significance (∼TFO & ∼TFR: r = 0.01, p-value = 0.61, N = 2032) maintaining its significance for the distance between the response and the last fixation (∼TFO & ∼TFR: *r* = 0.10, p-value < 10e-3, N=2032). This evidence suggests that participants who did not fixate on the target during the search were less consistent in their responses. However, those who correctly reported the target were able to gather information during the search that not only pointed to the location but also reduced the uncertainty of their decision.

### 3.2 Model Results

To model scanpaths in natural scenes we presented the nnELM model, an adaptation of the ELM model [Najemnik and Geisler, 2009]. For that, similar to the IBS model [Sclar et al., 2020, Bujia et al., 2022, Travi et al., 2022], we used CNNs to model the visual processing of image features. This model conveys slightly better results than previous models in the ViSioNS benchmark (see Supp. Material and Github). Furthermore, it runs up to 25 times faster than our previous best model (nnIBS), which allows us to better explore fixation-by-fixation metrics.

To compare participants’ manual responses with their previous fixations, we used the nnELM on the scanpaths (only fixations) and the extended scanpaths (all fixations plus the objective response as an additional fixation) (Fig. 4A). We first compared the performance of the nnELM in predicting the whole scanpaths using individual Human Scanpath Prediction (HSP) metrics [Travi et al., 2022], averaged across all fixations/responses (Fig. S2). Although significant, the changes in the distributions between scanpaths and the extended scanpaths were very small (Wilcoxon signed-rank, p-value < 10e-3, N= 7086; Fig. S2). This approach introduces only a small extension on the scanpath (1 extra fixation/response on an average of 6.15 previous fixations), thus a small effect size is expected. To solve this issue, we performed the analysis at the fixation level, we compared the prediction performance of the objective response against the previous fixations (Fig. 4 and Fig. S3). We aligned the fixations to the response, the 0 represents the last fixation, the -1 represents the penultimate, and so on, and we estimate individual HSP metrics for each fixation/response. There seemed to be a tendency for a better prediction for the responses in all metrics compared to any previous fixation independently of its order (Fig. 4D, E). We grouped all fixations and fit a linear mixed-effect model for each metric using as predictors the condition of the fixation and the trial condition (Table S2). As suggested in the previous analysis, the performance of the model in each of these metrics of the response location was better than any other fixation (AUC and IG in Fig. 4, t-value(isResp) = 8.55 and t-value(isResp) = 2.46 respectively; NSS and LL in Fig. S3, t-value(isResp) = 20.89 and t-value(isResp) = 6.98; Table S2). All main effects of the trial conditions (∼TFO & TFR and ∼TFO & ∼TFR) were notably negatively significant (Table S2) suggesting that the model had a higher agreement on trials where the subject found the target online. Yet, the strength of the effects seemed different according to the trial condition and metric. For instance, the AUC and the LL showed larger effects on the interaction terms for all conditions, but, on the contrary, not all interactions were significant for the IG metric.

**Figure 4:**
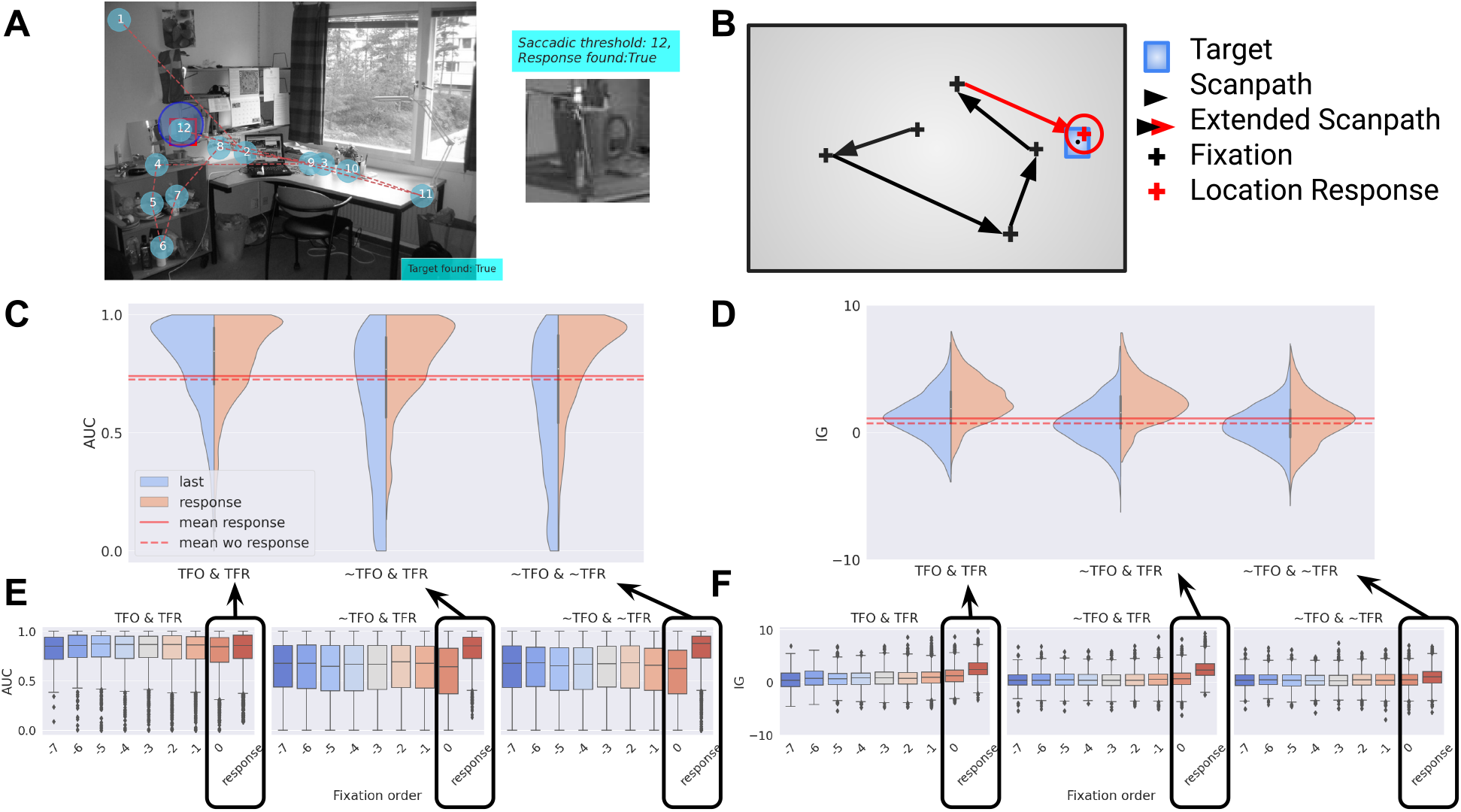
Predictive performance by behavior and category. **A**. Scanpath and objective response on the left side with the response marked with a blue circle where the center is the location response and the radius is the uncertainty response. On the right, there is an example of how we extended the scanpath. **B, C**. Distribution of AUC and IG for the last fixation and the response by trial type respectevely. Equivalent to the last two boxplots of panels A and B. The Red dashed line is the mean score over all fixations, red line is the mean over all fixations including objective responses. **D**. Distribution of AUC for the prediction of response-aligned fixations by trial type. **E**. Distribution of IG for the prediction of response-aligned fixations by trial type.

To determine the factors affecting the searcher’s uncertainty and the model’s effectiveness in prediction, we assessed whether the model could encode relevant information that correlates with the participant’s uncertainty. We hypothesized that the uncertainty relies on a few latent variables: how far the response was from the target or the last fixation, the extent of the image explored (scanpath length), and an intrinsic measure of the information gathered by the subject along the search (entropy or any topological feature). We built a series of LMMs with an incremental set of predictors to explain the uncertainty over the complete sample. Later, with the best model, we evaluated each trial outcome separately to identify how uncertainty was modulated by the behavior in each case.

Initially, we focused on the random effect structure (participants, images, or both) along with the distances to the target or the last fixation (Table S3), refining the model incrementally before evaluating the information-related variables in the subsequent step (Table S4). We did not include both distances in the same model because they were highly correlated (Pearson *r* = 0.51, p-value < 10e-3). The best model based on the AIC value was the one with the target-response distance as a fixed effect and both participants and images as random effects. In all cases, the distance effect was significant (t-values > 2, Table S3). Secondly, we incorporated the length of the scanpaths to the best of the previous models. This variable did also present a significant positive effect (t-value(ScanLen) = 3.22, Table S3) but the AIC augmented by 2 points. Despite the AIC getting slightly worse, we decided to include it considering that it is a relevant variable a priori, its individual effect resulted significant, and the conditional-R^2^ slightly improved.

Subsequently, we focused on evaluating the model-based information; we considered two distinct kinds of estimates of the 2D probability distribution: the entropy of the 2D decision map on which the model based the decision of where to perform the next fixation or the persistence homology of the same decision map. Regarding the 2D entropy, there are multiple proposals on how to measure it; we explored five approaches for calculating it: the Shannon entropy (**ent**), the fuzzy 2D entropy (**FEnt2D**), the permutation 2D entropy (**PEnt2D**), the sampling 2D entropy (**SEnt2D**), and distributional 2D entropy (**DEnt2D**).

The alternative approach proposed here is to quantify the information based on the topological data analysis using the H0 persistence homology of the model’s probability map. The hypothesis underlying this approach is that uncertainty may be influenced by the allocation of attention to a latent representation of the belief about the target’s location. For example, if this latent representation includes several potential sites where the target could be found after a few search steps, the uncertainty should be higher compared to cases where the probability mass is concentrated in a single dominant site. To test this, we computed for the decision map the 0-dimensional persistence diagram or barcode corresponding to the sublevel set filtration [Tralie et al., 2018] and calculated the number of connected components (number of bars) and the persistence entropy (see Methods). These last metrics provide a distinctive notion of information than classical entropies that could be complementary to them.

Only two of the considered variables resulted in significant results, the distributional 2D entropy (Table S4, t-val(DEnt2D) = 2.68, N=6883) and the persistence entropy (Table S4, t-val(PE) = -2.46, N=6883). The positive effect of the DEnt2D on the reported uncertainty suggests that as the spatial distribution of probability becomes more spread out or diverse, the subject perceives greater uncertainty. This indicates that when the model’s probability map showed multiple areas with significant probabilities, the subject felt less confident about the target’s location. On the other hand, since H0 persistence entropy measures the topological structure –specifically the number and lifetime of connected components–, a lower persistence entropy may indicate that there are fewer or less prominent topological features (fewer distinct regions where the probability is concentrated) indicating clearer, more defined clusters of high probability), the subject experiences less uncertainty. In such cases, the subject may be more confident (lower uncertainty) when the decision map has well-defined clusters of probable target locations. Both results suggest that the ELM is capturing relevant information that can encode aspects of the real uncertainty experienced by the subjects.

Finally, to understand the effect of those variables, we disaggregated the trials for the found condition (TFO vs ∼TFO), selected the best model previously built in terms of AIC (Fig. 5A) and fitted them separately. When looking at the cases where subjects found the target online (Fig. 5B), the impact of entropy diminished and lost its significance, while factors like scanpath length and target-response distance persisted. This suggests that participants were less affected by entropy-based information once they found the target, and they were efficiently assessing the distance to the target with their response size. In contrast, when subjects did not find the target, the entropy effect was significant (Fig. 5C) but both the intercept and the scanpath length effect turned out to be not significant. This implies that in cases where the target was missed, the model effectively could capture some of the uncertainty of the participants with higher uncertainty in cases where the entropy was bigger. However, since that data contained participants that later on found (or not) the target with their response, we fitted an additional model with a categorical variable for Target Found Response condition (∼TFO & TFR, Fig. 5D). The categorical variable had a negative impact on the response size as we would have expected from previous results (Table S1), meaning that subjects that found the target with their response did it confidently. What resulted more interesting is that even segmenting the effect of entropy remained significant, increasing the uncertainty, meaning that in all cases the model’s decision map may be effectively encoding part of the underlying uncertainty of the searcher.

**Figure 5:**
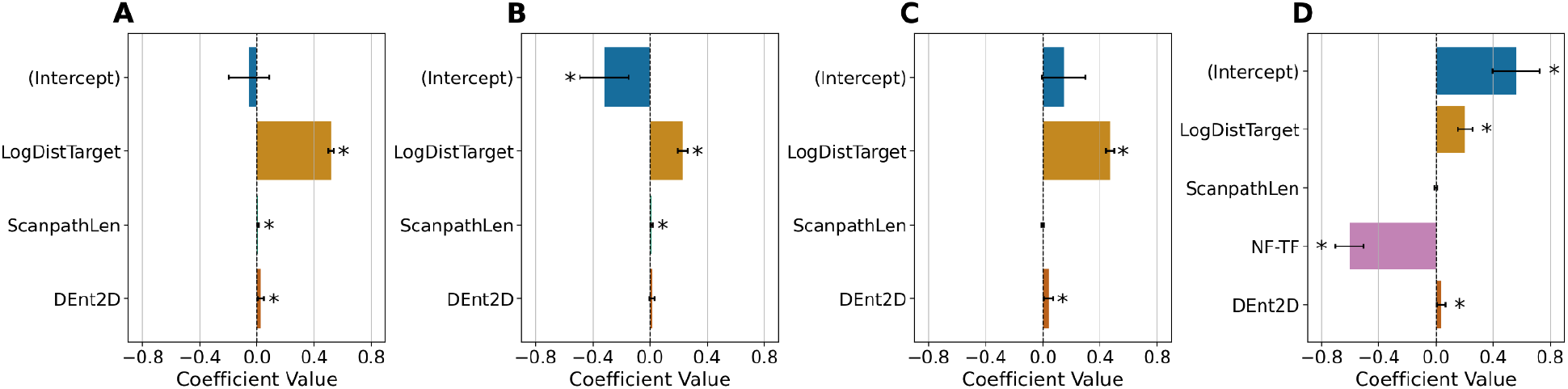
Linear mixed-effects models on the estimation of the uncertainty. The dependent variable was the uncertainty estimated as the response size. We evaluated image id (img) and participant id (subj) as random effects. As fixed effects, we considered: the distance measured as the Euclidean distance from the response to the target (in pixels); scanpath length (scanL, in fixations); and an entropy measure: Dent2D = distributional 2D entropy. Confidence intervals were estimated using profiling, and an asterisk indicated whether the absolute t-value was larger than 1.96. **A**. uses all trials, and **B**. uses trials in which the target was fixated during the search (TFO). **C**. and **D**. use trials in which the target was not fixated during the search (∼TFO) and the last one (**D**) includes a logical variable indicating whether it was correctly reported in the explicit response or not (∼TFO & TFR).

## 4 Discussion

Understanding how individuals navigate a scene while searching for an object provides a unique lens to study decisionmaking processes. This process, often studied in artificial images, gains two distinct advantages when investigated in naturalistic scenes. First, the use of everyday life stimuli—whether sounds, images, movies, or texts—offers greater ecological validity. Second, naturalistic tasks are intrinsically sequential, involving a complex interplay of perceptual, cognitive, and motor processes that must be integrated to accomplish the task. In our experiment, participants engaged in a crowded scene search that required iteratively fixing patches, analyzing them, deciding on the presence of the target, integrating information, selecting the next location, and shifting attention, a complex dynamic and layered decision-making process.

Building on previous studies that primarily focused on comparing human and model scanpaths or predicting the location of the next fixation, this work addresses a relatively underexplored aspect: uncertainty during visual search in naturalistic settings. We do this by integrating uncertainty reports, eye movement data, and computational tools. Crucially, our use of the adapted ELM model revealed that uncertainty is a multifaceted process influenced by factors such as the length of the scanpath, the distance between the perceived and the true target location, and the entropy of probabilistic decision maps.

The ELM model emerged as the top-performing approach in the ViSioNS benchmark [Travi et al., 2022], demonstrating a strong capability to predict human fixations and variations in uncertainty. Unlike prior models, the ELM captures response locations and predicts uncertainty during visual search. By linking spatial probability distributions and information features of decision maps to subjective uncertainty, we provide evidence that confidence reflects not only accuracy but also more nuanced belief structures. For instance, scenarios with high uncertainty (e.g., trials where the target was not found) revealed significant correlations between participants’ uncertainty responses and the model’s internal states.

These findings extend beyond theoretical significance to practical applications. The ability to model and predict uncertainty during visual search has implications for fields such as AI, human-computer interaction, and neuropsychology. For example, understanding how uncertainty evolves during decision-making can inform the design of adaptive systems in diagnostic imaging, where accurate visual search is critical. Similarly, incorporating uncertainty metrics into AI models could enhance their interpretability and reliability in dynamic environments, such as autonomous vehicles or surveillance systems.

While our study provides valuable insights, several methodological considerations warrant attention. First, the use of a single task type, the nature of the stimuli (grayscale images) and the specific computational model limits the generalizability of our findings. Future research should explore the applicability of the ELM model across diverse tasks and stimuli to test its robustness. Second, the reliance on pre-trained saliency maps may introduce biases unrelated to human perceptual processes. Addressing these limitations will strengthen the broader applicability of the proposed framework.

Our work opens several avenues for future research. A promising direction is the integration of neural measures, such as EEG or fMRI, with computational outputs to investigate the neural correlates of choice and uncertainty. For instance, EEG could be used to examine how neural activity reflects transitions between fixations and uncertainty in real-time. Additionally, further adaptations of the ELM model could include exploring the role of task complexity or extending the framework to multi-modal stimuli. These directions would provide a deeper understanding of the relationship between internal representations and subjective uncertainty.

In conclusion, this study provides a paradigm for decoding the latent cognitive processes underlying visual search behavior by combining behavioral data with computational modeling. Our findings emphasize the importance of uncertainty as a key aspect of decision-making and highlight the potential of advanced computational models to capture its nuanced dynamics.

## Acknowledgments

We thank P. Lagomarsino and J. Laurino for their collaboration with the data acquisition, and Fermin Travi for collaborating in the development of the ViSioNS Benchmark from which we use the HSP metrics. We would also like to thank Matias J. Ison for critical suggestions and feedback. P. Lagomarsino was an undergraduate student in the lab during the early phase of this project. His untimely passing is a great loss to all who knew him. He approached every task with boundless enthusiasm and an unwavering motivation to learn, inspiring everyone around him. We will always remember and cherish his passion and spirit.

## Author contributions

MS, GS and JK designed the task, collected human data, and defined the model idea. MS prepared the indoor images/targets dataset. GB performed the modeling effort. GR contributed to the implementation of the model. GB and JK performed the analysis and wrote the manuscript. All authors reviewed the manuscript, contributed to the article and approved the submitted version.

## Funding information

The authors were supported by the CONICET and the University of Buenos Aires (UBA). The data collection was supported by the CONICET (PIP 11220150100787CO), the ARL (Award W911NF1920240), and the National Agency of Promotion of Science and Technology (PICT 2018-2699).

## Data and code availability

The code and data to replicate the results obtained in this manuscript can be found in Github’s repository. This code also uses functions available in the Visions Benchmark to estimate human scanpath prediction metrics and to compare to other state-of-the-art models.

## Competing Interests

The authors declare no conflict of interest.

## Notes

### Competing Interest Statement

The authors have declared no competing interest.

