## Supporting material for "Uncertainty during visual search: Insights from a computational model and behavioral experiment in natural stimuli"

PREPRINT SUBMITTED TO BIORXIV

**Gaston Bujia[1,2], Gonzalo Ruarte[1,2], Melanie Sclar[3], Guillermo Solovey[4], and Juan E. Kamienkowski[*,2,5,*]**

[1]**Laboratorio de Inteligencia Artificial Aplicada (LIAA), Instituto en Ciencias de la Computación (ICC), CONICET - Universidad de Buenos Aires. Buenos Aires, Argentina.**
[2]**Departamento de Ciencias de la Computación, Facultad de Ciencias Exactas y Naturales, Universidad de Buenos Aires. Buenos Aires, Argentina.**
[3]**University of Washington, Seattle, Washington, United States.**
[4]**Instituto de Cálculo, Universidad de Buenos Aires-CONICET, Buenos Aires, Argentina.**
[5]**Maestría en Explotación de Datos y Descubrimiento del Conocimiento, Facultad de Ciencias Exactas y Naturales, Universidad de Buenos Aires. Buenos Aires, Argentina.**
[*]******

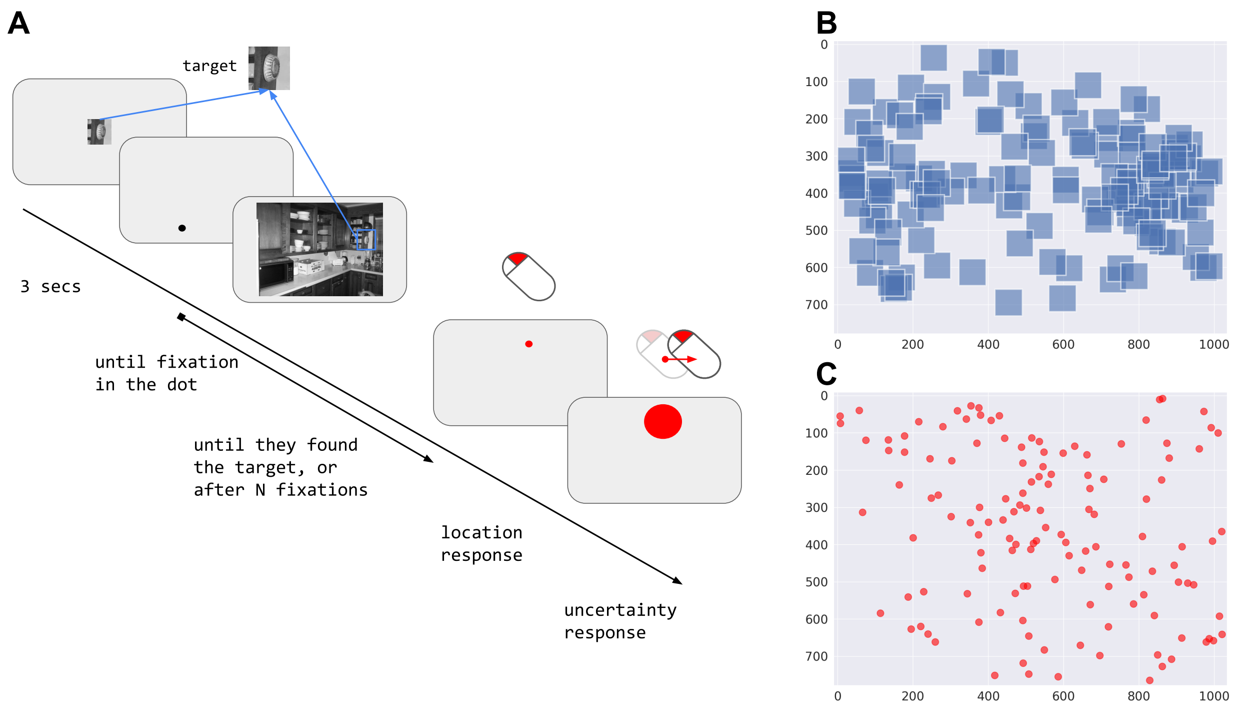


Figure 1: **Experimental design. A.** Timeline. The target was displayed at the center of the screen on a gray background at twice its original size for 3 seconds. After that, a dot was presented until fixated, starting the trial. The trial finished when the target was found or after the participant performed N fixations. After the trial ended, the participant reported the target location by selecting a point with the left mouse button and then reported their uncertainty by determining the radius length using the mouse buttons. **B.** Spatial distribution of targets (blue squares), and **C.** Spatial distribution of first fixations (red dots).

$$p_i(T+1) = \frac{p_i(T)\exp\left[d^2_{ik(T+1)}W_{ik(T+1)}\right]}{\sum_{j=1}^{n} p_j(T)\exp\left[d^2_{jk(T+1)}W_{jk(T+1)}\right]} \tag{1}$$

This can be rewritten more generally as:

$$p_i(T) = \frac{\text{prior}(i)\prod_{t=1}^{T}\exp\left(d^2_{ik(t)}W_{ik(t)}\right)}{\sum_{j=1}^{n}\text{prior}(j)\prod_{t=1}^{T}\exp\left(d^2_{jk(t)}W_{jk(t)}\right)} \tag{2}$$

The visibility map $d_{ik(T)}$ represents how visible the target is on the location $i$ when the observer is located at $k(T)$ (it is quantified as a fixed 2D Gaussian centered on the current fixation). Meanwhile, $W_{ik(T)}$ represents the template response that measures the information collected from the location $i$ given the observer location $k(T)$. This template response was originally a random sample drawn from a Gaussian distribution $W_{ik(T+1)} \sim N(\mu_{ik(T)}, \sigma_{ik(T)})$ that depended only on the presence or absence of the target and the inverse of the visibility map (eq. 3). Here, we incorporate the notion of similarity between the target and each image location (i.e. similarity map, see below)

$$\mu_{ik(t)} = \mathbb{I}(i = \text{target location}) - 0.5\ ; \quad \sigma_{ik(t)} = \frac{1}{d_{ik(t)}} \tag{3}$$

To calculate the next fixation, the ELM estimates the expected reduction of entropy as the current posterior probability distribution across the possible target locations weighted by the visibility (eq. 4):

$$E[\Delta H(T+1)|k(T+1)] = \frac{1}{2}\sum_{i=1}^{n} p_i(T)d^2_{ik(T+1)} \tag{4}$$

Given a fixation order $T$, eq. 4 inherently defines a mapping between all possible target locations on the scene and the expected information gain for each of them. This map is used later for calculating the human scanpath metrics (see Metrics section) and we call it decision map. Finally, the next fixation $k$ is selected as the one that maximizes the information gain obtained from selecting that location $k_{opt}(T+1)$ (eq. 5):

$$k_{\text{opt}}(T+1) = \arg\max_{k(T+1)} \left\{ \sum_{i=1}^{n} p_i(T) d_{ik(T+1)}^2 \right\} \tag{5}$$

The original ELM approach was entirely proposed, and tested in artificial stimuli where subjects had to search for a Gabor patch embedded in 1/f noise. Further adaptations were proposed for modeling other characteristics of the scanpaths like the fixation duration [Zhou and Yu, 2021] or incorporating color [Rashidi et al., 2023]. In the case of natural stimuli, we followed the original adaptation proposal made in Bujia et al. [2022], Sclar et al. [2020], Travi et al. [2022], the idea is to introduce a notion of similarity map $\phi$ into the template response via modifications into the parameters of the Gaussian distribution. Previously, it was performed with a mean value that combined the visibility and the similarity calculated with cross-correlation or structural similarity between the target template and the i-th location of the image ($\phi_i$) [Sclar et al., 2020, Bujia et al., 2022] and, later on, with the attention map from a deep neural network [Zhang et al., 2018]. We will use the output of the IVSN network [Zhang et al., 2018] at location *i* (eq. 7) in combination with the proposal for the template response (eq. 6):

$$\widetilde{W}_{ik(t)} \sim \mathcal{N}(\widetilde{\mu}_{ik(t)}, \widetilde{\sigma}_{ik(t)}^2) \tag{6}$$

$$\widetilde{\mu}_{ik(t)} = \mu_{ik(t)} \cdot \left( d_{ik(t)} + \frac{1}{2} \right) + \phi_i \cdot \left( \frac{3}{2} - d_{ik(t)} \right) \tag{7}$$

$$\widetilde{\sigma}_{ik(t)} = \frac{1}{a \cdot d_{ik(t)} + b} \tag{8}$$

Table 1: Classification of trials according to the search and response results. TFO: target found online (during the search) and TFR: target found response (with the manual response after the search).

| | **TFR** | **~TFR** |
|---|---|---|
| **TFO** | 45.52% (3322) | 5.99% (437) |
| **~TFO** | 23.35% (1704) | 25.14% (1835) |

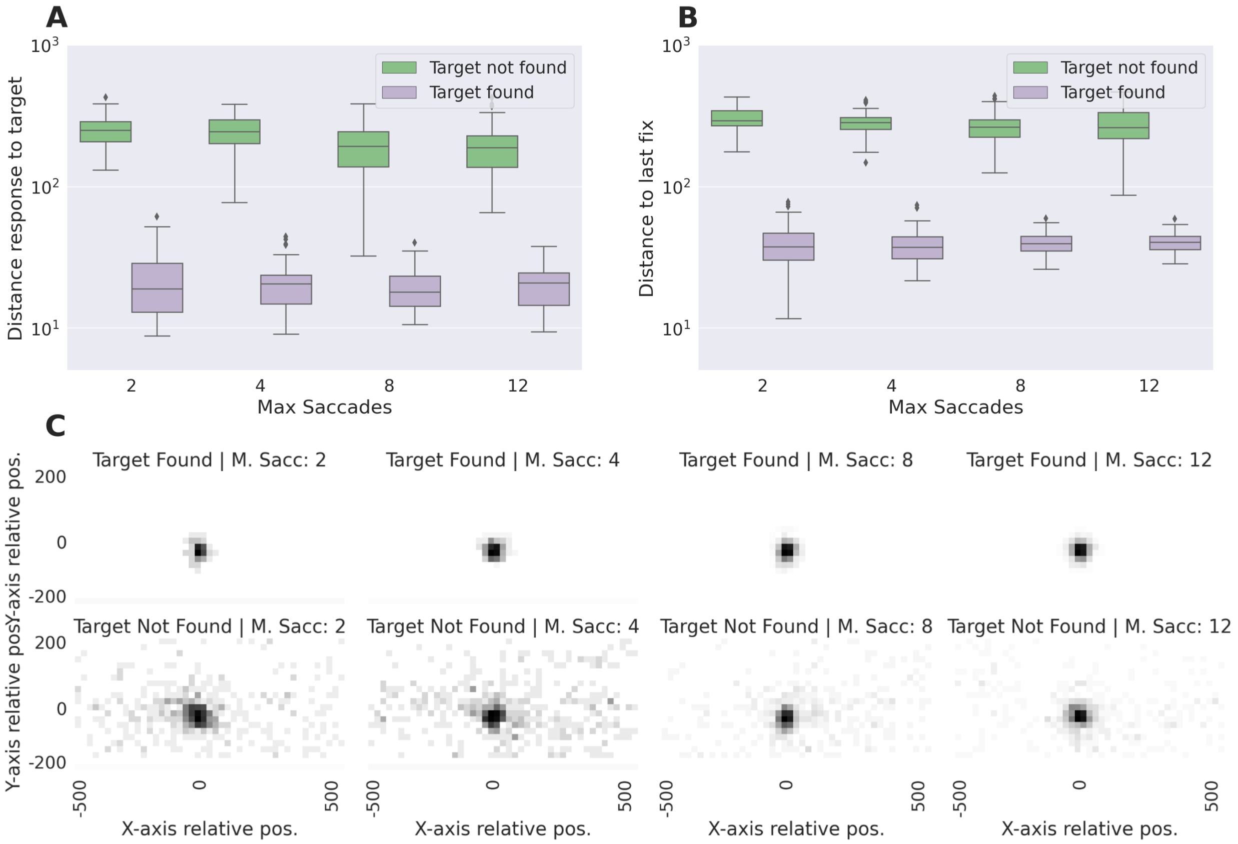


Figure 2: **Objective response. A.** Euclidean distance from the response to the target, **B.** Euclidean distance from the response to the last fixation, and, C. spatial distribution of responses relative to the target at the center, all as a function of the found condition of the trial (found or not found) and the maximum number of saccades allowed. (Each point is subject, error bars correspond to the standard deviation). In the rightmost of panel **C.**, the color bar represents the density mapping for all the heatmaps in that row.

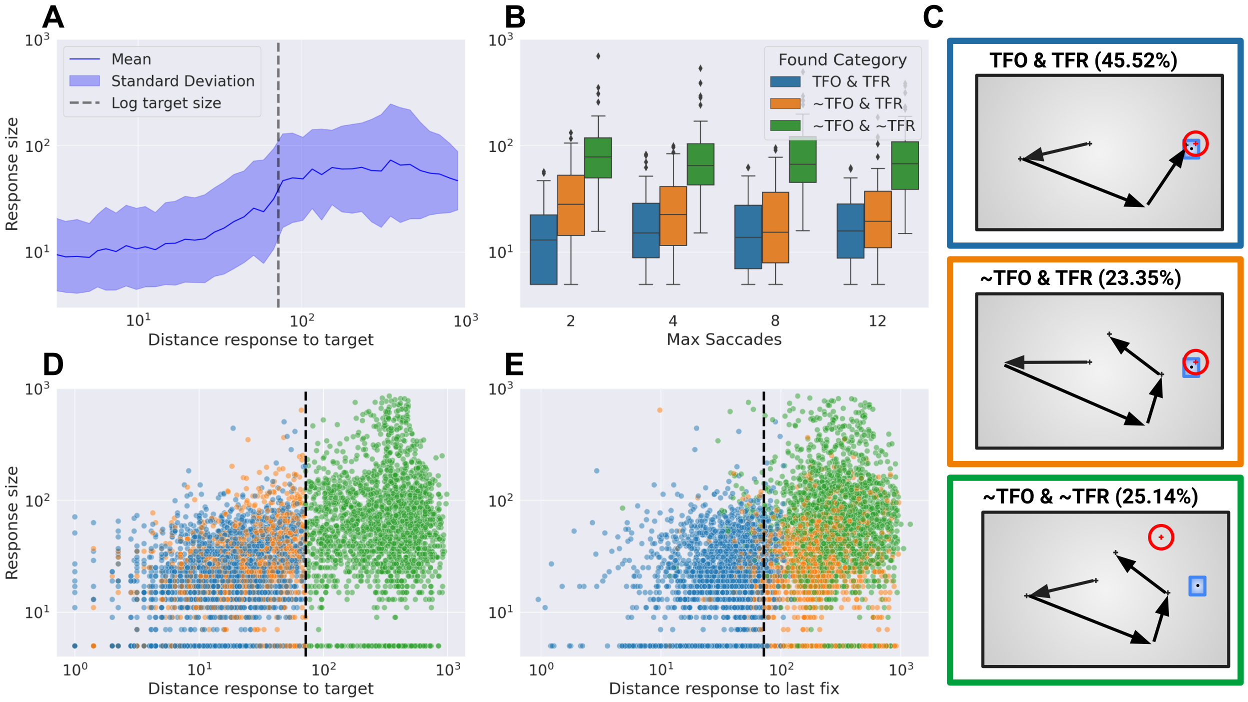


Figure 3: **Uncertainty response.** **A.** Logarithm of the uncertainty (or response size) as a function of the distance of the response to the target for all responses considered. **B.** Uncertainty, measured as the size of the response, as a function of the maximum of saccades allowed and grouped by the trial category. **C.** Scheme of the trial categories. **D.** Distribution of distance from response to target and the response size for each trial category and distribution of distance from last subject fixation to the response and the response size (**E.**). The dashed line represents the threshold for considering a response as target found.

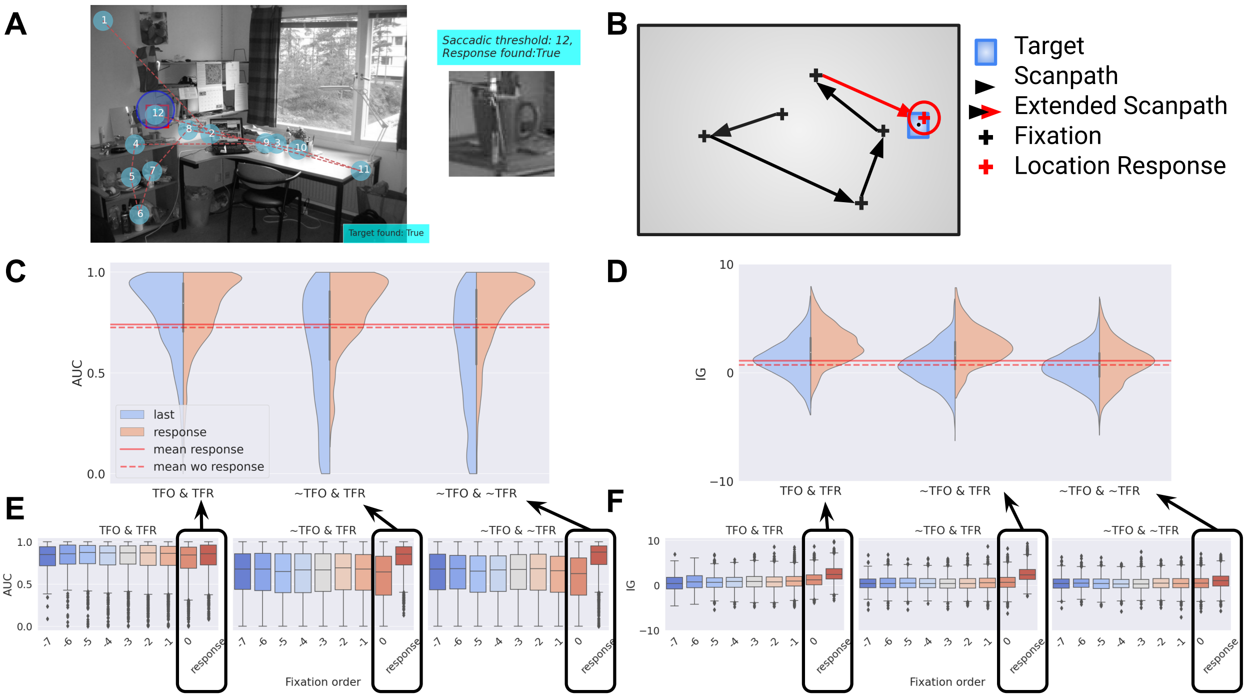


Figure 4: **Predictive performance by behavior and category. A.** Scanpath and objective response on the left side with the response marked with a blue circle where the center is the location response and the radius is the uncertainty response. On the right, there is an example of how we extended the scanpath. **B**, **C.** Distribution of AUC and IG for the last fixation and the response by trial type respectevely. Equivalent to the last two boxplots of panels A and B. The Red dashed line is the mean score over all fixations, red line is the mean over all fixations including objective responses. **D.** Distribution of AUC for the prediction of response-aligned fixations by trial type. **E.** Distribution of IG for the prediction of response-aligned fixations by trial type.

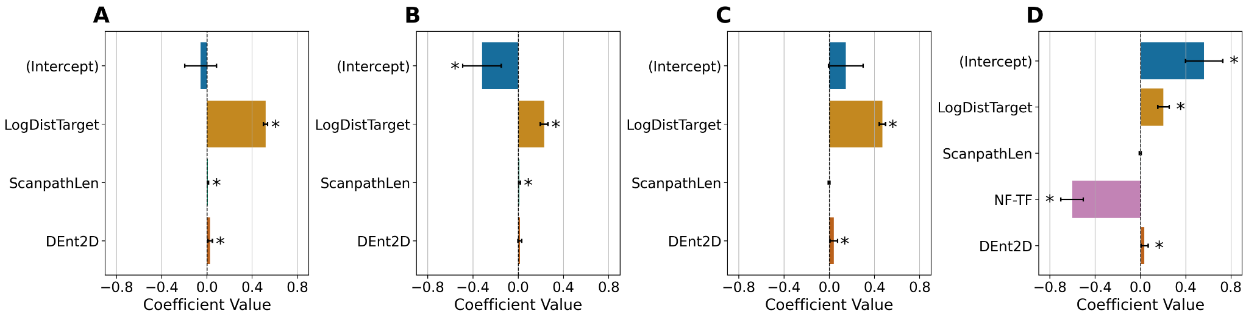


Figure 5: **Linear mixed-effects models on the estimation of the uncertainty.** The dependent variable was the uncertainty estimated as the response size. We evaluated image id (img) and participant id (subj) as random effects. As fixed effects, we considered: the distance measured as the Euclidean distance from the response to the target (in pixels); scanpath length (scanL, in fixations); and an entropy measure: Dent2D = distributional 2D entropy. Confidence intervals were estimated using profiling, and an asterisk indicated whether the absolute t-value was larger than 1.96. **A.** uses all trials, and **B.** uses trials in which the target was fixated during the search (TFO). **C.** and **D.** use trials in which the target was not fixated during the search (~TFO) and the last one (**D**) includes a logical variable indicating whether it was correctly reported in the explicit response or not (~TFO & TFR).

#### Competing Interests

The authors declare no conflict of interest.

### Supplemental Materials

#### Supplemental Figures

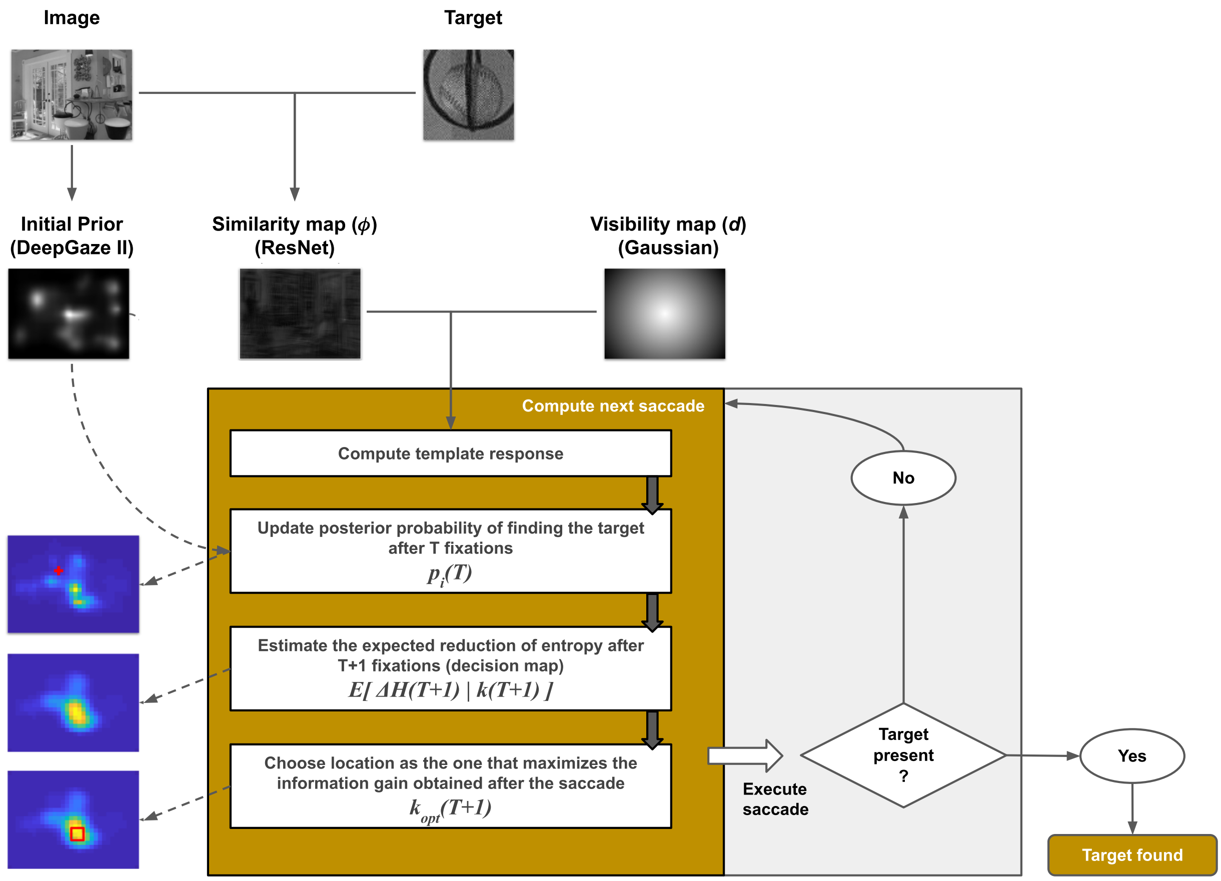


Figure S1: **Visual Search Model: Workflow of the nnELM model**. The nnELM model receives the search image and the target (exact object or category) as inputs, also it can use the first fixation location as input. DeepGaze II model generates a prior distribution of the first fixation if not provided. Then, this prior and a fixed visibility map (Gaussian distribution centered on the current fixation) along with a target similarity map derived from a ResNet model [He et al., 2015, Zhang et al., 2018], are used to calculate the likelihood of possible fixation points. In subsequent fixation cycles, the prior is updated from the previous cycle's posterior. The current fixation posterior is then computed using this likelihood and the prior. A decision mechanism, either Bayesian [Sclar et al., 2020, Bujia et al., 2022] or ELM, uses this posterior to determine the next fixation position. This process repeats iteratively, with each cycle's posterior becoming the prior for the next cycle until the target is identified or the maximum amount of fixations is met.

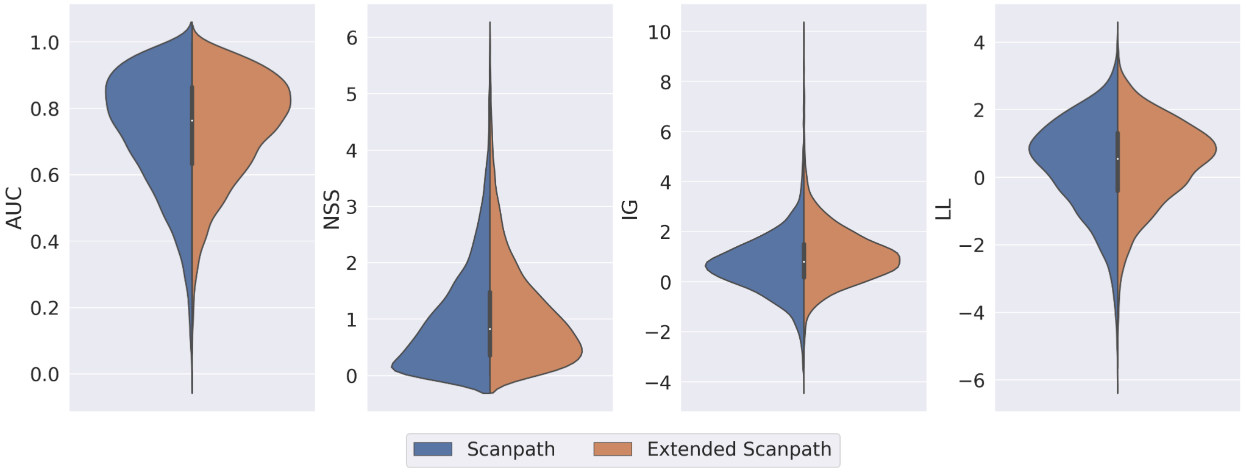


Figure S2: **Full scanpath vs. extended scanpath on the 4 metrics.**

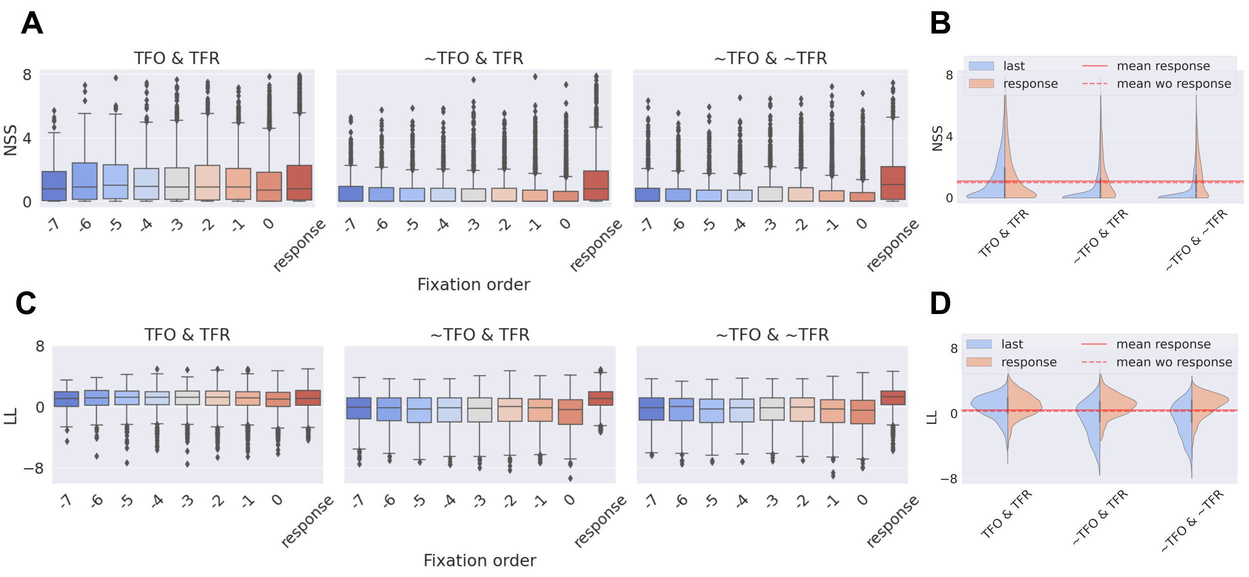


Figure S3: **Response vs. previous fixations.** **(A-B)** NSS and **(C-D)** LL metric for each of the trial categories comparing the distribution of the previous seven fixations to the response location. The last column **(B, D)** corresponds specifically to the NSS and LL metric for each trial category, comparing the distribution only of the last fixation against the response location.

#### Supplemental Tables

Table S1: **Linear mixed-effects models of the distance from the response to the target (log) and the last fixation (log)**. Continuous variables were normalized to have 0 mean and deviation equal to 1. Variables were transformed using the natural logarithm. Reported CIs correspond to the 95% confidence. MaxFix is the maximum allowed number of saccades (plus one); TF corresponds to Target Found trials; ~TFO & TFR corresponds to Target Not Found Online and Target Found on Response; ~TFO & ~TFR corresponds to Target Not Found Online and Target Not Found on Response.

| ID | Target Variable | Variable | Coefficient (95% CI) | t-value (95% CI) |
|---|---|---|---|---|
| 1 | LogDistTarget | (Intercept) | 0.73 (0.71, 0.75) | **62.75** (60.79, 64.71) |
| | | MaxFix | -0.01 (-0.01, -0.01) | **-11.42** (-13.39, -9.46) |
| | | TF | -0.26 (-0.29, -0.24) | **-21.36** (-23.33, -19.40) |
| | | MaxFix:TF | 0.01 (0.00, 0.01) | **4.76** (2.80, 6.72) |
| 2 | LogDistLastFix | (Intercept) | 0.79 (0.78, 0.80) | **168.58** (166.62, 170.55) |
| | | MaxFix | -0.00 (-0.01, -0.00) | **-9.00** (-10.99, -7.02) |
| | | TF | -0.29 (-0.30, -0.27) | **-40.25** (-42.23, -38.28) |
| | | MaxFix:TF | 0.00 (0.00, 0.01) | **5.97** (4.01, 7.93) |
| 3 | LogRespSize | (Intercept) | 3.80 (3.61, 4.00) | **38.99** (37.02, 40.96) |
| | | MaxFix | -0.04 (-0.05, -0.03) | **-8.53** (-10.49, -6.57) |
| | | TF | -1.11 (-1.24, -0.98) | **-16.63** (-18.60, -14.67) |
| | | MaxFix:TF | 0.03 (0.02, 0.05) | **5.22** (3.26, 7.19) |
| 4 | LogRespSize | (Intercept) | 2.55 (2.35, 2.74) | **25.65** (23.69, 27.62) |
| | | MaxFix | 0.00 (-0.01, 0.01) | 0.17 (-1.79, 2.13) |
| | | ~TFO & ~TFR | 1.55 (1.43, 1.68) | **23.90** (21.93, 25.88) |
| | | ~TFO & TFR | 0.50 (0.36, 0.63) | **7.28** (5.32, 9.24) |
| | | MaxFix:~TFO & ~TFR | -0.01 (-0.02, 0.00) | -1.76 (-3.72, 0.20) |
| | | MaxFix:~TFO & TFR | -0.03 (-0.04, -0.01) | **-3.91** (-5.88, -1.95) |

Table S2: **Linear mixed-effects models on the estimation of the visual search metrics according to the behavior**. AUC was transformed with logit and the NSS was transformed with the natural logarithm. *isResp* is a binary variable that indicates whether the data point considered is a response or a fixation; ~TFO & TFR corresponds to Target Not Found Online and Target Found on Response; ~TFO & ~TFR corresponds to Target Not Found Online and Target Not Found on Response.

| ID | Target Variable | Variable | Coefficient (95% CI) | t-value (95% CI) |
|---|---|---|---|---|
| 1 | LogitAUC | (Intercept) | 1.80 (1.65, 1.95) | **23.11** (21.14, 25.07) |
| | | ~TFO & ~TFR | -1.29 (-1.39, -1.20) | **-26.72** (-28.68, -24.76) |
| | | ~TFO & TFR | -1.41 (-1.51, -1.32) | **-28.32** (-30.28, -26.35) |
| | | isResp | 0.33 (0.26, 0.41) | **8.55** (6.59, 10.51) |
| | | ~TFO & ~TFR:isResp | 1.33 (1.21, 1.45) | **21.47** (19.51, 23.43) |
| | | ~TFO & TFR:isResp | 1.31 (1.18, 1.44) | **19.12** (17.16, 21.08) |
| 2 | LogNSS | (Intercept) | -0.12 (-0.23, -0.01) | **-2.11** (-4.08, -0.15) |
| | | ~TFO & ~TFR | -0.35 (-0.45, -0.26) | **-7.04** (-9.00, -5.08) |
| | | ~TFO & TFR | -0.45 (-0.55, -0.34) | **-8.54** (-10.50, -6.58) |
| | | isResp | 0.08 (0.02, 0.15) | **2.46** (0.50, 4.42) |
| | | ~TFO & ~TFR:isResp | 0.52 (0.40, 0.63) | **8.72** (6.76, 10.68) |
| | | ~TFO & TFR:isResp | 0.35 (0.22, 0.48) | **5.30** (3.34, 7.26) |
| 3 | IG | (Intercept) | 1.37 (1.16, 1.59) | **12.42** (10.46, 14.38) |
| | | ~TFO & ~TFR | -0.72 (-0.89, -0.56) | **-8.52** (-10.48, -6.56) |
| | | ~TFO & TFR | -0.59 (-0.76, -0.41) | **-6.65** (-8.61, -4.69) |
| | | isResp | 1.44 (1.30, 1.57) | **20.89** (18.93, 22.85) |
| | | ~TFO & ~TFR:isResp | -0.89 (-1.11, -0.68) | **-8.12** (-10.08, -6.16) |
| | | ~TFO & TFR:isResp | 0.23 (-0.01, 0.47) | 1.90 (-0.06, 3.86) |
| 4 | LL | (Intercept) | 0.74 (0.60, 0.89) | **9.89** (7.93, 11.86) |
| | | ~TFO & ~TFR | -1.45 (-1.54, -1.36) | **-31.05** (-33.01, -29.09) |
| | | ~TFO & TFR | -1.56 (-1.65, -1.46) | **-32.24** (-34.21, -30.28) |
| | | isResp | 0.26 (0.19, 0.33) | **6.98** (5.02, 8.94) |
| | | ~TFO & ~TFR:isResp | 1.56 (1.45, 1.68) | **26.16** (24.20, 28.12) |
| | | ~TFO & TFR:isResp | 1.51 (1.38, 1.64) | **22.81** (20.85, 24.77) |

Table S3: **Linear mixed-effects models on the estimation of the uncertainty based on behavioral variables.** The dependent variable was the uncertainty estimated as the response size over the complete dataset. We evaluated image id (img) and participant id (subj) as random effects. As fixed effects, we considered: the distance measured as `dist_last_fix` = distance from the response to the last fixation (in pixels) or `dist_target` = distance from the response to the target (in pixels); scanpath length (scanL, in fixations). The Akaike Information Criterion (AIC) and the conditional pseudo-$R^2$ were used to compare between models. Supporting data with additional information may be found in the Github repository.

| **Model** | **Variable** | **t-value** | **AIC** | **Pseudo-$R^2$ Condicional** |
|---|---|---|---|---|
| `dist_last_fix + (1|img)` | Distance<br>ScanLen | 32.74<br>- | 17853 | 0.206 |
| `dist_target + (1|img)` | Distance<br>ScanLen | 51.00<br>- | 16811 | 0.322 |
| `dist_target + (1|subj)` | Distance<br>ScanLen | 49.22<br>- | 15499 | 0.500 |
| `dist_last_fix + (1|subj)` | Distance<br>ScanLen | 69.24<br>- | 15010 | 0.500 |
| `dist_target + (1|img) + (1|subj)` | Distance<br>ScanLen | 58.31<br>- | 13758 | **0.578** |
| `dist_target + ScanLen + (1|img) + (1|subj)` | Distance<br>ScanLen | 57.03<br>3.22 | 13760 | **0.579** |

Table S4: **Linear mixed-effects models on the estimation of the uncertainty including entropy measures.** The dependent variable was the uncertainty estimated as the response size. We evaluated image id (`img`) and participant id (`subj`) as random effects. As fixed effects, we considered: the distance measured as `dist_target` = distance from the response to the target (in pixels); the scanpath length (`scanL`, in fixations); and an entropy measure: `Ent` = Shannon entropy, `FEnt2D` = fuzzy 2D entropy, `PEnt2D` = permutation 2D entropy, `SEnt2D` = sampling 2D entropy, or `DEnt2D` = distributional 2D entropy. Supporting data with additional information may be found in Github's repository.

| Model | Distance (t-value) | Scanpath (t-value) | Entropy (t-value) | AIC | Conditional Pseudo-$R^2$ |
|---|---|---|---|---|---|
| `dist_target + ScanLen + H0f + (1\|img) + (1\|subj)` | 56.93 | 3.22 | 0.04 | 13769 | 0.579 |
| `dist_target + ScanLen + PE + (1\|img) + (1\|subj)` | 56.83 | 3.17 | -2.46 | 13768 | 0.580 |
| `dist_target + ScanLen + H0 + (1\|img) + (1\|subj)` | 56.83 | 3.13 | 0.00 | 13767 | 0.580 |
| `dist_target + ScanLen + PEf + (1\|img) + (1\|subj)` | 56.92 | 3.17 | 0.05 | 13767 | 0.580 |
| `dist_target + ScanLen + Ent + (1\|img) + (1\|subj)` | 56.99 | 3.24 | 0.03 | 13765 | 0.580 |
| `dist_target + ScanLen + SEnt2D + (1\|img) + (1\|subj)` | 57.02 | 3.23 | 0.08 | 13765 | 0.579 |
| `dist_target + ScanLen + FEnt2D + (1\|img) + (1\|subj)` | 56.96 | 3.27 | 0.17 | 13768 | 0.580 |
| `dist_target + ScanLen + PEnt2D + (1\|img) + (1\|subj)` | 56.98 | 3.15 | -1.11 | 13768 | 0.580 |
| `dist_target + ScanLen + DEnt2D + (1\|img) + (1\|subj)` | 56.80 | 3.18 | -2.68 | 13762 | 0.580 |
